# Interactions among Diameter, Myelination and the Na/K pump Affect Axonal Resilience to High Frequency Spiking

**DOI:** 10.1101/2021.03.25.437049

**Authors:** Yunliang Zang, Eve Marder

**Affiliations:** Volen Center and Department of Biology, Brandeis University, Waltham, MA 02454, USA

**Keywords:** computational model, action potential, node of Ranvier, electrogenic pumps, sodium dynamics

## Abstract

Axons reliably conduct action potentials between neurons and/or other targets. Axons have widely variable diameters and can be myelinated or unmyelinated. Although the effect of these factors on propagation speed is well studied, how they constrain axonal resilience to high frequency spiking is incompletely understood. Maximal firing frequencies range from ~ 1 Hz to > 300 Hz across neurons, but the process by which Na/K pumps counteract Na^+^ influx is slow, and it is unclear the extent to which slow Na^+^ removal is compatible with high frequency spiking. Modeling the process of Na^+^ removal shows that large diameter axons are more resilient to high frequency spikes than small diameter axons, because of their slow Na^+^ accumulation. In myelinated axons, the myelinated compartments between nodes of Ranvier act as a ‘reservoir’ to slow Na^+^ accumulation and increase the reliability of axonal propagation. We now find that slowing the activation of K^+^ current can increase the Na^+^ influx rate, and the effect of minimizing the overlap between Na^+^- and K^+^-currents on spike propagation resilience depends on complex interactions among diameter, myelination and the Na/K pump density. Our results suggest that, in neurons with different channel gating kinetic parameters, different strategies may be required to improve the reliability of axonal propagation.

**Significance Statement:** The reliability of spike propagation in axons is determined by complex interactions among ionic currents, ion pumps and morphological properties. We use compartment-based modeling to reveal that interactions of diameter, myelination and the Na/K pump determine the reliability of high frequency spike propagation. By acting as a ‘reservoir’ of nodal Na^+^ influx, myelinated compartments efficiently increase propagation reliability. Although spike broadening was thought to oppose fast spiking, its effect on spike propagation is complicated, depending on the balance of Na^+^ channel inactivation gate recovery, Na^+^ influx and axial charge. Our findings suggest that slow Na^+^ removal influences axonal resilience to high frequency spike propagation, and that different strategies may be required to overcome this constraint in different neurons.

## Introduction

Axons usually reliably conduct information (spikes) to other neurons, muscles and glands. The largest diameter axons are found in some invertebrates, where the squid giant axon and arthropod giant fibers are parts of rapid escape systems. In the mammalian nervous system, axon diameters can differ by a factor of > 100. Some are covered with myelin sheaths but others are not (1). Both myelination and large axon diameter increase spike propagation speed (2, 3), and the latter factor can also increase axonal resilience to noise perturbation (4).

Neuronal firing frequency is not static, but variations in firing rates are used to encode a wide range of signals such as inputs from sensory organs and command signals from motor cortex. The maximal firing frequency, critical for neuronal functional capacity, ranges from ~ 1 Hz to > 300 Hz (5–8). However, it remains unclear how axonal physical properties constrain axonal resilience to high frequency firing. During action potentials, Na^+^ flows into axons and then ATP is used by Na/K pumps to pump out the excess Na^+^. Given the limited energy supply in the brain, evolutionary strategies including shortening spike propagation distance, sparse coding and reducing Na^+^- and K^+^-current overlap etc. have helped minimize energy expenditure (9–13). However, the observation that high frequency spiking tends to occur in large diameter axons (5) is confusing because fast spiking in large diameter axons causes more Na^+^ influx and exerts a heavy burden on the energy supply in the brain (14, 15). Additionally, the process by which Na/K pumps remove Na^+^ is slow (16–22). In neurons, it takes seconds or even minutes to remove the excess Na^+^ after [Na^+^] elevation (16–21). It remains unknown how these directly correlated processes, fast spiking and slow Na^+^ removal, interact in the control of reliable spike generation and propagation. Furthermore, it is unclear whether reducing Na^+^- and K^+^-current overlap can consistently decrease the rate of Na^+^ influx and accordingly enhance axonal reliability to propagate high frequency spikes. Consequently, by modeling the process of Na^+^ removal in unmyelinated and myelinated axons, we systematically explored the effects of diameter, myelination and Na/K pump density on spike propagation reliability.

## Results

### Different [Na^+^] Decay Dynamics in Unmyelinated and Myelinated Axons

We implemented multicompartment models of axons in NEURON (23). Both the unmyelinated and myelinated axon models include I_Na_, I_KD_, I_KA_, I_leak_ and the Na/K pump. All channels and pumps were evenly spread in the unmyelinated axon model and at the nodes of Ranvier in the myelinated axon model. For the myelinated compartments, there is a significantly lower density of I_leak_ and C_m_ to simulate the effect of insulation (Materials and Methods). The model structures are shown in Fig. 1A.

**Figure 1.**
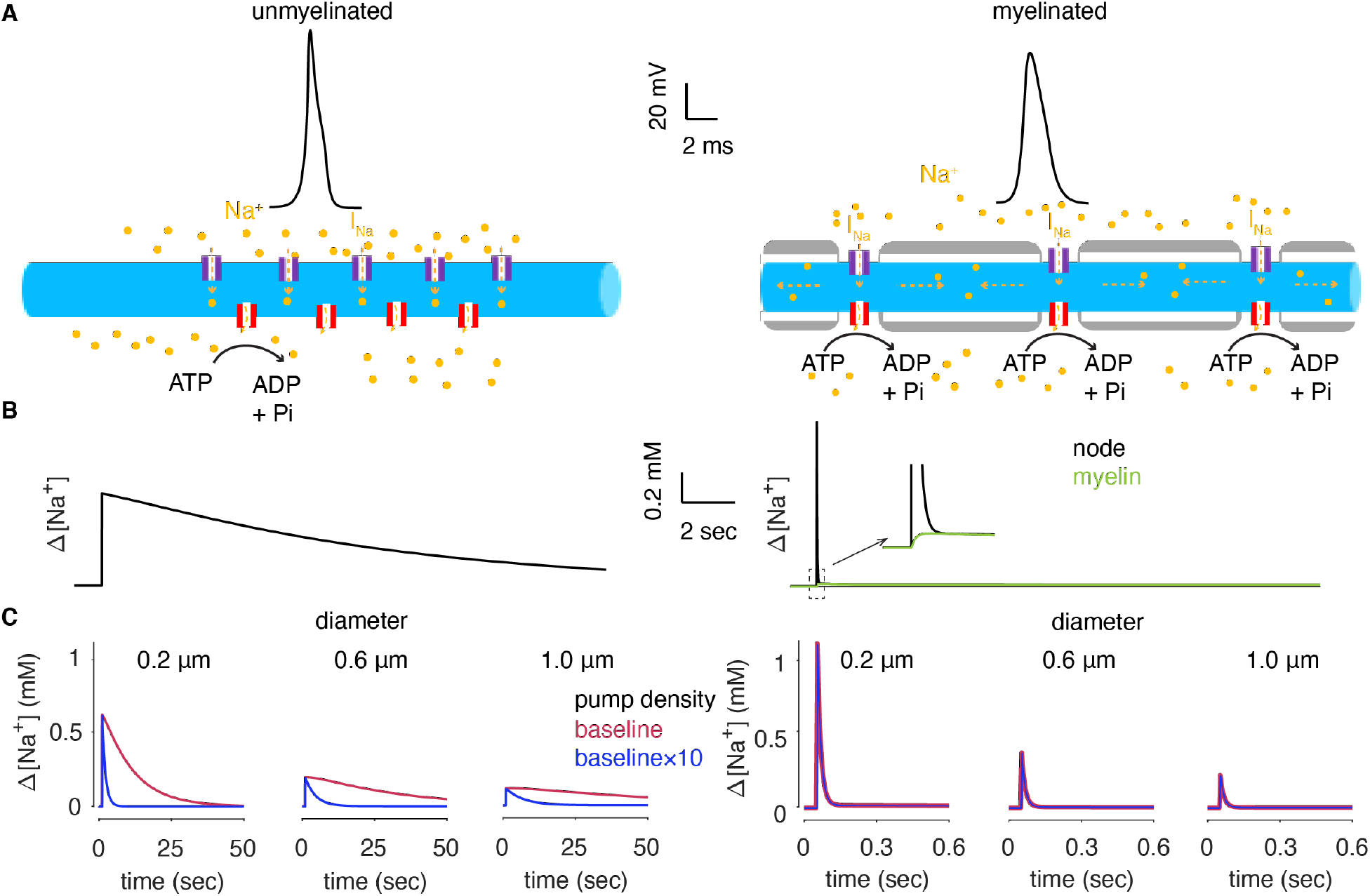
Spike-triggered [Na^+^] rise and decay in unmyelinated and myelinated axon models. (A) Schematics of the unmyelinated (left) and the myelinated (right) axon models. Black traces show spike waveforms in two models (50 μm from the distal axon end; In myelinated model, it is also a node of Ranvier). (B) Spike-triggered [Na^+^] rise and decay in the unmyelinated (left) and the myelinated (right) axon models. Black traces were measured at corresponding voltage recording sites. Green trace shows [Na^+^] in the myelinated compartment (16.7 μm from the node of Ranvier) and the initial part is enlarged in the inset. The Na/K pump density is 0.5 pmol/cm^2^, referred to as baseline condition hereafter. (C) The effects of diameter and pump density on [Na^+^] rise and decay in the unmyelinated (left) and the myelinated (right, at the node of Ranvier) axon models. Note the very different time scales in the two models.

Spikes propagate rapidly in both models. In the unmyelinated axon model, the propagation increases from 0.1 m/s to 0.3 m/s when the axonal diameter increases from 0.2 μm to 1 μm. In the myelinated axon model, the speed increases from 0.87 m/s to 2.1 m/s accordingly. Upon the arrival of spikes, Na^+^ channels open and extracellular Na^+^ flows into axons to raise the intracellular [Na^+^] immediately (Fig. 1B). Then K^+^ currents are activated to repolarize the spikes. Compared with the other ionic currents, the magnitude of the Na/K pump current is extremely small during a single spike, but it decays very slowly. Eventually, the Na/K pump gradually removes Na^+^ from the axon, leading to a slow, many second, [Na^+^] decay in the unmyelinated axon model (Fig. 1B, left). In contrast, in the myelinated axon model, the spike-triggered [Na^+^] elevation at the node of Ranvier decays on the time scale of hundreds of ms (Fig. 1B, right). The Na/K pump densities in the two models are the same and thus the quick decay is not a result of super-efficient pumps. In the myelinated axon model, Na^+^ channels are placed only in the nodes of Ranvier and there is no transmembrane Na^+^ influx in the myelinated compartments. But, the slightly elevated [Na^+^] in the myelinated compartment and the rapid decay of nodal [Na^+^] are due to the axial diffusion of Na^+^ into neighboring myelinated compartments, rather than immediate pump action.

Because of the large diffusion coefficient (24) and the lack of buffering (16), [Na^+^] in the radial direction becomes rapidly homogeneous. Thus, spike-triggered [Na^+^] elevation can be well predicted by the surface-to-volume ratio (SVR ∝ 1/diameter). In both models, spike-triggered [Na^+^] elevation decreases with diameter (Fig. 1C). Increasing the pump density speeds up the [Na^+^] decay in the unmyelinated axon model, but has no obvious effect on the nodal [Na^+^] decay in the myelinated axon model, further confirming that the quick decay is not a result of the Na/K pump.

### Na^+^ Accumulation in the Unmyelinated Axon at High Firing Frequencies

Compared with the basal level before spiking (~ 10 mM), the [Na^+^] elevation caused by a single spike is small and insufficient to affect axonal excitability (Fig. 1B). However, neurons can fire at high frequencies in certain behavioral contexts (7, 25–27). With continuous high frequency spiking, Na^+^ can accumulate if subsequent spikes occur before the complete recovery of a preceding spike elicited [Na^+^] rise. In the unmyelinated axon model, Na^+^ significantly accumulates within just 100 spikes, and the accumulation speed increases with firing frequency because of a faster Na^+^ influx rate (Fig. 2A). Larger SVRs can also make Na^+^ accumulate faster in thin axons. Note that the seemingly slower Na^+^ accumulation in the 0.2-μm-diameter axon at 60 Hz is due to earlier propagation failure. In parallel with Na^+^ accumulation, the Na^+^ equilibrium potential (driving force of Na^+^ current) becomes more negative and the action potential peak decreases (Fig. 2B, C). With increased firing frequency, propagation starts to fail. The failure starts from 30 Hz in the 0.2-μm-diameter axon (Fig. 2C), but from 60 Hz in larger diameter axons. A repertoire of propagation failure patterns were observed. In Fig. 2D1, every other spike propagated through the axon, but a more complicated propagation failure pattern is shown in Fig. 2D2.

**Figure 2.**
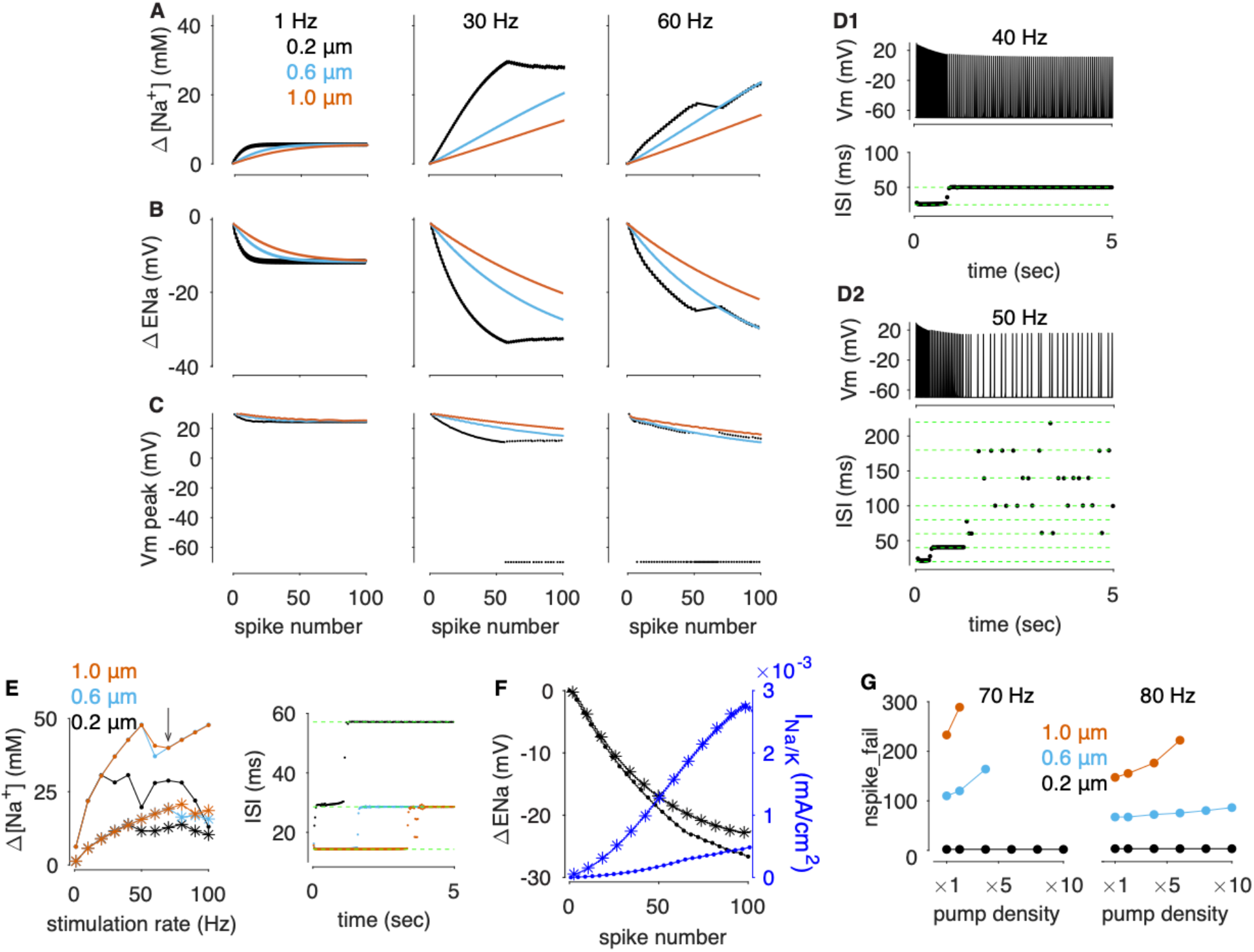
The effects of diameter and pump density on Na^+^ accumulation and resilience to high frequency spiking in the unmyelinated axon model. A-C show [Na^+^] increase, Na^+^ equilibrium potential decrease and membrane potential peak reduction (bottom) in models with different diameters and firing frequencies. (D) Voltage traces and interspike intervals (ISIs) at distal axon for 40 Hz (D1) and 50 Hz (D2) spike signals in the 0.2-μm-diameter axon. (E) Steady state [Na^+^] elevation at different stimulation frequencies (left). Right panel shows ISIs at distal axon for 70 Hz spike signals with baseline pump density. (F) Na^+^ equilibrium potential decrease (black) and outward pump current increase (blue) with different pump densities. 80 Hz spike signals in the 0.6-μm-diameter axon. In E & F, dot and star represent 1 and 10 times of baseline pump density, respectively. (G) The effect of pump density on axonal resilience to high frequency spiking, measured by the number of spikes required to cause propagation failure (nspike_fail), when firing at 70 and 80 Hz. The absence of data points means propagation doesn’t fail under corresponding conditions.

When the model fires over a sufficiently long period, the Na^+^ influx rate and efflux rate gradually balance each other to make [Na^+^] reach a steady state. Despite different accumulation speeds, the steady state [Na^+^] increases with firing frequency and is independent of diameter until propagation starts to fail (Fig. 2E). The rate of Na^+^ influx per surface area is only determined by the firing frequency because of identical Na^+^ channel density. The Na/K pump-caused net Na^+^ efflux rate is determined by −3*k*_1_[*pump*][*Na*^+^]^3^ + 3*k*_2_[*pumpNa*]. Because the pump density is also identical, the model needs identical [Na^+^] to match influx rate and reach a steady state irrespective of diameters. If propagation fails, the steady state [Na^+^] depends on the actual pattern of spikes that propagate through the axon. For example, at 70 Hz with baseline pump density, the steady state [Na^+^] is the same in the 0.6- and 1.0-μm-diameter axons but lower in the 0.2-μm-diameter axon, because of their specific propagation failure patterns (Fig. 2E). When pumps remove excess Na^+^, they also carry a negative current, which gradually repolarizes the membrane potential during spikes along with Na^+^ accumulation. The factors causing the gradual occurrence of failure includes the Na^+^ channel’s incomplete recovery from inactivation, decreased driving force due to Na^+^ accumulation, and increased outward pump current during continuous spiking. This propagation failure can be eliminated or alleviated when Na^+^ accumulation is not considered in the model. Increasing pump density consistently reduces Na^+^ accumulation (Fig. 2E) to alleviate the equilibrium potential decrease, but also enhances the outward pump current to directly decrease the axonal excitability (Fig. 2F).

Na^+^ accumulates slowly in large diameter axons and in these axons it always requires more spikes to trigger propagation failure than in small diameter axons, making these axons more resilient to high frequency firing (Fig. 2G). At higher frequencies, Na^+^ accumulates faster and Na^+^ channels recover less from inactivation. Consequently, fewer spikes are required to trigger propagation failure and accordingly the resilience decreases (Fig. 2G). With higher pump densities in the model, the axonal resilience to high frequency spiking is increased (Fig. 2 G), suggesting that the alleviated decrease in Na^+^ equilibrium potential prevails against the enhanced outward pump current (Fig. 2F).

### Myelinated Compartments Slow Na^+^ Accumulation

In contrast to the slow decay of [Na^+^] in the unmyelinated axon, the [Na^+^] at the node of Ranvier in the myelinated axon shows a quick decay (Fig. 1B). Can nodal [Na^+^] elevate at high firing frequencies? It does elevate to reduce the Na^+^ equilibrium potential and action potential peak in parallel (Fig. 3A-C). As explained previously, the quick decay of nodal [Na^+^] is a result of nodal Na^+^ quickly diffusing into neighboring myelinated compartments rather than being removed. After diffusion, the nodal [Na^+^] is still higher than the pre-spiking level (Fig. 1B). In other words, nodal Na^+^ influx not only needs to fill the nodes of Ranvier, but also the neighboring myelinated compartments, which is equivalent to decreasing the SVR. Consequently, the nodal Na^+^ still accumulates, but at a much slower rate (Fig. 3A, F). It requires thousands of spikes to significantly change nodal [Na^+^] in myelinated axons. Na^+^ in the myelinated compartments flows back during the removal of Na^+^ by nodal Na/K pumps. Therefore, by acting like an ‘anti-flood water reservoir’, myelinated compartments slow down the nodal Na^+^ accumulation and the excitability reduction.

**Figure 3.**
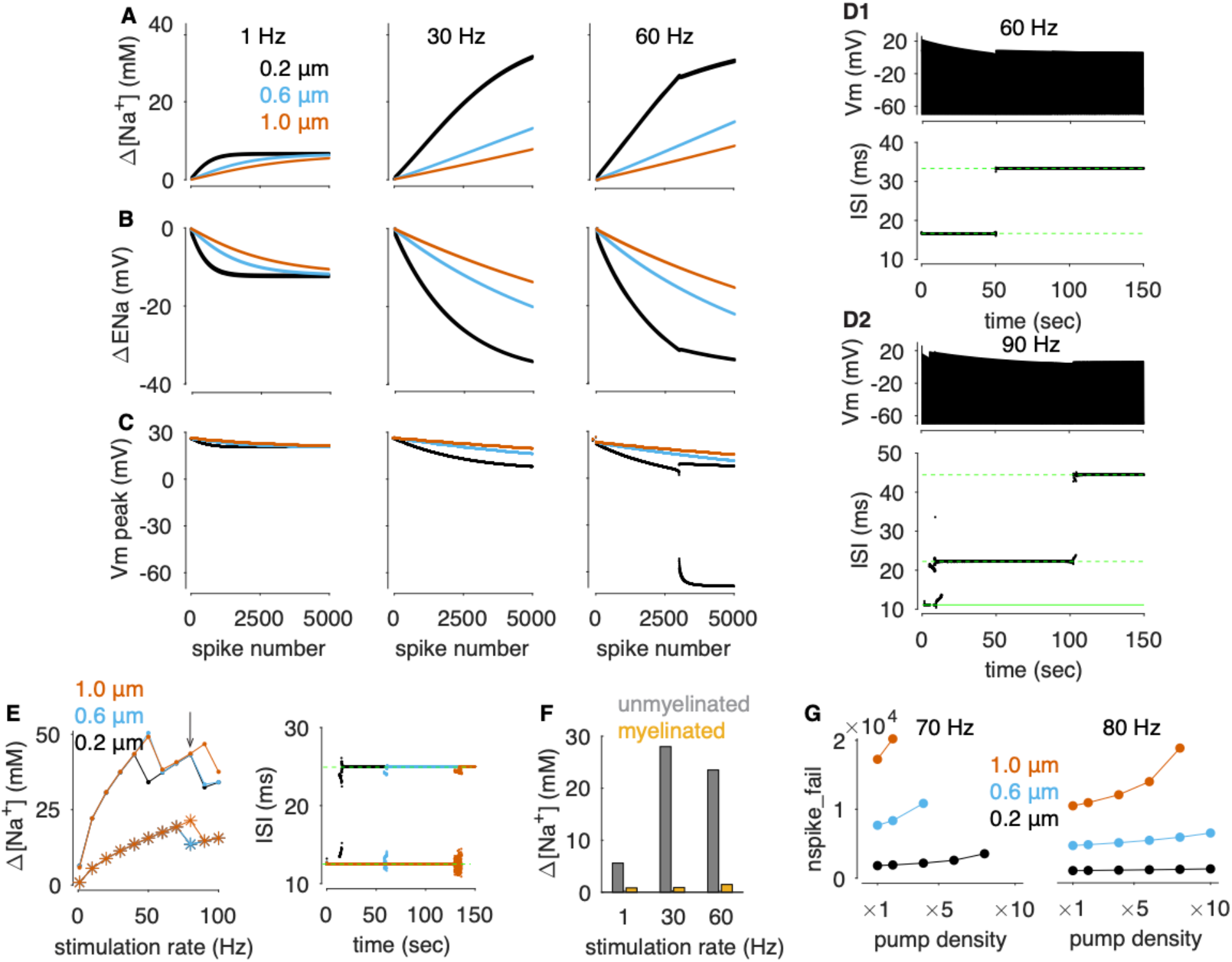
Myelination slows Na^+^ accumulation to increase axonal resilience to high frequency spiking. A-C show [Na^+^] increase, Na^+^ equilibrium potential decrease and membrane potential peak reduction at the node of Ranvier in the myelinated axon model with different diameters and firing frequencies. (D) Voltage traces and ISIs at distal axon for 60 Hz (D1) and 90 Hz (D2) spike signals in the 0.2-μm-diameter axon. (E) Steady state [Na^+^] elevation at different stimulation frequencies (left). Dot and star represent 1 and 10 times of baseline pump density, respectively. Right panel shows ISIs at distal axon for 80 Hz spike signals with baseline pump density. (F) 100-spike caused Na^+^ accumulation in the unmyelinated axon and the myelinated axon (node of Ranvier). The diameter is 0.2 μm, with baseline pump density. (G) The effect of pump density on axonal resilience to high frequency spiking, when firing at 70 Hz and 80 Hz. The absence of data points means propagation doesn’t fail under corresponding conditions.

Similar to the case in unmyelinated axon models, nodal Na^+^ accumulates faster at higher firing frequencies (Fig, 3A). Propagation failure starts from 50 Hz in the 0.2-μm-diameter axon, but from 60 Hz in larger diameter axons. In Fig. 3D1, every other spike can propagate through the axon. However, only every fourth spike can successfully propagate in Fig. 3D2. The steady state [Na^+^] is diameter independent until propagation fails at high frequencies (Fig. 3E). When firing at 80 Hz with baseline pump density, the steady state [Na^+^] is still the same in axons with different diameters, because they eventually have the same propagation patterns.

Because Na^+^ accumulates more slowly in myelinated axons compared with unmyelinated axons (Fig. 3F), the propagation is more reliable in myelinated axons (Fig. 3G). Large diameter axons are consistently more resilient and the resilience decreases with firing frequency. Similar to unmyelinated axon models, more pumps increase the propagation resilience.

### The Effects of Na^+^-and K^+^-Current Overlap on Propagation Resilience

Previous work has shown that the temporal overlap between the Na^+^- and K^+^-currents strongly affects Na^+^ influx during spiking (8). Can minimizing their temporal overlap consistently enhance axonal resilience by reducing the rates of Na^+^ influx and Na^+^ accumulation?

Slowing the activation of I_KD_ was reported to increase spike duration and decrease Na^+^ influx per spike (8). However, slowing I_KD_ activation increases spike triggered Na^+^ influx instead of decreasing it in our model (Fig. 4A, c→d). By systematically varying I_KD_ activation time constant in the range of 0.2-1.4 times of the baseline value, we observed a U-shaped relationship between Na^+^ influx rate and I_KD_ activation rate. Slowing the activation rate initially decreases the Na^+^ influx rate, but further slowing it increases the Na^+^ influx rate (Fig. 4A). This occurs because slowing I_KD_ activation increases spike duration to widen the Na^+^ current profile; on the other hand, slower repolarization reduces the Na^+^ channel driving force to decrease the current amplitude during the repolarization phase. Their balance determines the net change of Na^+^ influx rate caused by slowing the I_KD_ activation.

**Figure 4.**
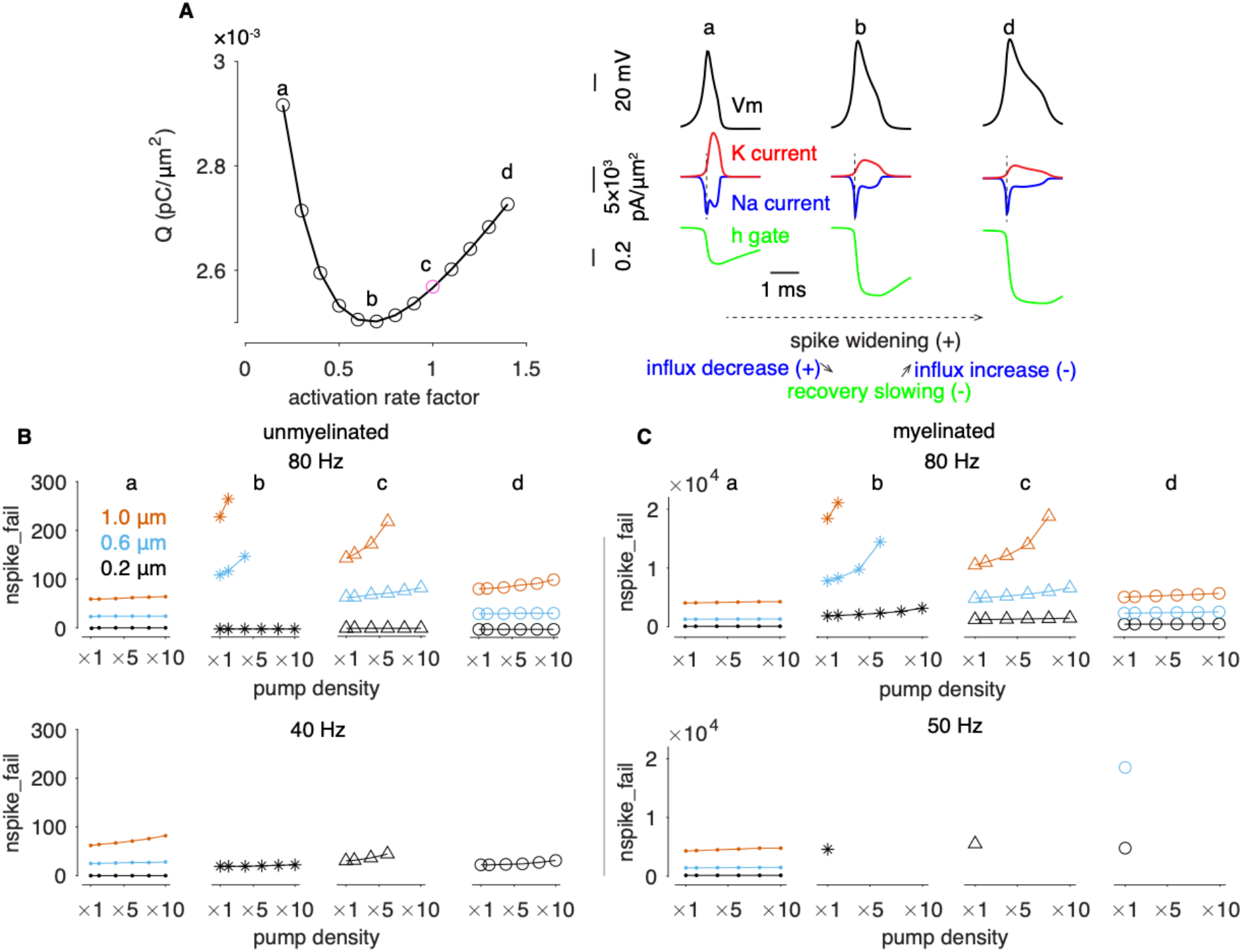
Complex effects of slowing I_KD_ activation on Na^+^ influx rate and propagation resilience. (A) U-shaped dependence between Na^+^ influx rate and I_KD_ activation rate. y axis (left panel) measures single spike triggered Na^+^ influx per μm^2^. c represents the baseline model (magenta); a, b and d represent 0.2, 0.6 and 1.4 times of the baseline I_KD_ activation time constant, respectively. Changes of spike shape, current profile and Na^+^ channel inactivation gate state with slowed I_KD_ activation. The effects of delayed activation of I_KD_ on propagation resilience in unmyelinated and myelinated models are shown in B and C, respectively. a - d correspond to conditions defined in A. Resilience changes are measured by comparing symbols of the same color (coding diameter). For example, from a to b at 40 Hz in the unmyelinated model, black stars are higher than black dots, suggesting increased resilience in the 0.2-μm-diameter axon. The resilience also increases in larger diameter axons, because slowed I_KD_ activation eliminates propagation failure. The absence of data points means propagation doesn’t fail under corresponding conditions.

Apart from the U-shaped Na^+^ influx rate changes, slowing I_KD_ activation consistently increases spike duration, which results in changes that both favor and oppose spike propagation. Broader spikes provide more axial charge to neighboring segments per spike to favor spike propagation, but they also cause larger Na^+^ channel inactivation and less recovery to oppose spiking. These factors interact to make the effect of slowing I_KD_ activation on propagation reliability complex (Fig. 4B, C). In both unmyelinated and myelinated axon models, slowing I_KD_ activation from a to b consistently increases the propagation resilience. However, when slowing I_KD_ activation from b to c, the resilience increases at low firing rates but decreases at high firing rates in both models. Slowing I_KD_ activation from c to d consistently reduces the resilience in both models. These complex effects reflect the interactions among Na^+^ influx rate, spike duration and Na^+^ channel inactivation gate state changes.

We also explored the effect of speeding Na^+^ current inactivation on propagation reliability. Faster Na^+^ current inactivation consistently reduces spike duration and Na^+^ influx, but it also has complex effects on the resilience because of the interactions among the above-mentioned opposing factors (Fig. S1). The effects of diameter and myelination on the speed of Na^+^ accumulation and propagation resilience hold in all the simulations we have done.

## Discussion

Neuronal firing frequency often varies over a broad range to encode a wide range of signals. For a given neuron, the rate of Na^+^ entry at its high firing frequency is usually much higher than the rate at which excess Na^+^ is pumped out of the cell. Therefore, while short periods of high frequency firing should present no problem for most neurons, continuous high frequency spiking could result in compromised function, as intracellular [Na^+^] increases. Hence, we explore the dynamic relationships between fast spiking and slow Na^+^ removal for reliable spike generation and propagation.

The importance of the Na/K ATPase in regulating neuronal activities has attracted attention in recent years (28). It can set neuronal excitability and contribute to maintaining synchronous epileptiform bursting in developing CA3 neurons (29). By interacting with h-current, it regulates bursting activity in the leech heartbeat central pattern generator (30). There have also been theoretical efforts to study the pump’s role in controlling the transition between different firing patterns (31–33) and the transition between normal and pathological brain states (34–37). However, to the best of our knowledge, how the pump interacts with axon diameter and myelin to remove Na^+^ influx and affect axonal excitability has not been previously explored.

Axonal morphology can exert a critical constraint on a neuron’s maximal firing frequency. Large diameter and myelinated axons are intrinsically advantageous to propagate spikes because of their larger space constants. In this work, we highlight another advantage in propagating continuous high frequency spikes. Lower SVRs and slower Na^+^ accumulation make them more resilient to high frequency spiking. This may explain why fast-spiking parvalbumin interneurons are extensively myelinated (38) and why high frequency spiking tends to occur in large diameter axons, despite a higher energy cost of information transmission (5, 14, 15). Note that rapid redistribution of ions such as Na^+^ can occur at other sites and in other processes. For example, the soma, acting as a Na^+^ sink, can enhance reliable spike initiation in axon initial segments (16). Redistribution of chloride in dendrites is also shown to be important for chloride equilibrium potential and synaptic inhibition (39).

Increasing the Na/K pump density to remove Na^+^ more efficiently is expected to facilitate axonal resilience to high frequency spiking. Existing data argue that Na^+^ removal after spiking is slow. In axon initial segments of L5 pyramidal neurons, as a result of Na^+^ diffusing into the soma and myelin wrapped compartments, elevated [Na^+^] requires several seconds to approximately recover (16). In mitral cell tuft dendrites, after a short period of high frequency stimulation, it takes minutes for the Na/K ATPase to pump out the accumulated Na^+^ (17). This slow Na^+^ removal may be functionally important because it enables integration and binding of multiple chemosensory stimuli over a prolonged time scale (40). We used pump densities that should fall into a reasonable biological range, and the Na^+^ accumulation at high firing frequencies shown here is consistent with recent Na^+^ imaging data (17, 20).

This work illustrates the limitations of using conventional frequency-current curves generated with a short-term current injection to estimate the reliability of spike propagation (41, 42). Neither of these measures reflects the accommodation of neuronal excitability after continuous firing caused changes such as Na^+^ (explored in this work) and Ca^2+^ accumulation (showing a biphasic decay (43, 44)).

In the models explored here, myelinated axons are resilient to high frequency spiking elicited Na^+^ accumulation. Given that myelinated axons are usually much thicker (1, 45), spike propagation in myelinated axons should be extremely robust to fast spiking caused Na^+^ accumulation under usual physiological conditions. However, under conditions requiring neurons to fire for extended time at high frequencies, Na^+^ accumulation may still compromise the function of axons. Of course, the frequency threshold and the number of spikes required to cause spike failure will depend on the specifics of the expression of neuronal conductances, and will likely show variability as a consequence of variable channel densities (46). Additionally, intrinsic channel noise can cause spike failure (4, 27), and the normal function of the Na/K ATPases requires ATP, the supply of which has a limit (35). Both a shortage of ATP after continuous firing and channel noise can further reduce axonal reliability of propagating spikes.

In recent years, a trade-off has been proposed between a neuron’s capability to spike rapidly and the high energetic cost of spiking (8, 47). To achieve high spike rates (narrow spikes), neurons need to consume more energy to remove the larger Na^+^ influx, coproduct of fast spiking. However, in our simulations, we find a more complicated relationship between Na^+^ influx and spike duration (Fig. 4). Furthermore, although narrow spikes can facilitate high frequency spike generation, they are more vulnerable to Na^+^ accumulation or noise disturbance during propagation. This work implies that different strategies may be required in different types of neurons to guarantee reliable spike propagation for their functional requirements.

## Materials and Methods

### Axonal Morphology

We modeled a segment of axon with the length of 3131 μm. The model was implemented in NEURON (23). The model code is available from ModelDB (accession number: 267009). The simulations used an integration step of 0.05 ms. The unmyelinated model was discretized into 51 segments in the longitudinal direction. For the myelinated axon model, there are 31 nodes of Ranvier flanked with 32 myelinated compartments (Figure 1). The lengths of the nodes of Ranvier are 1 μm (nseg = 1). Each myelinated compartment is 100 μm long (nseg = 3), except the outer two myelinated compartments are 50 μm each (nseg = 3). In the radial direction, there are 4 shells to model Na^+^ diffusion (23). The Na^+^ diffusion coefficient is 0.6 μm^2^/ms (24) and there is no buffering (16), leading to a homogeneous [Na^+^] in the radial direction. In both peripheral and central nervous systems, unmyelinated axons are usually thinner than myelinated axons. In the central nervous system, axons > 0.2 μm are usually myelinated (1). To delineate the effect of both diameter and myelination on Na^+^ accumulation, we chose the same range of diameters between 0.2–1.0 μm.

### Ionic Channels and the Na/K pump

The model has a minimal set of ionic channels: fast Na current (I_Na_), delayed rectifier K current (I_KD_), A-type K current (I_KA_) and leak current (I_leak_). They have standard kinetic formulations in the form of 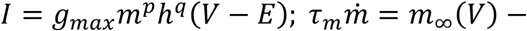 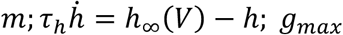 represents maximal conductance; *m* and *h* represent activation and inactivation gates; *p* and *q* represent corresponding gate exponents; *E* represents equilibrium potential; *τ*_*m*_ and *τ*_*h*_ represent corresponding time constants; All the channel kinetics were modified from (48). For I_Na_, *g*_*max*_ = 0.015 *S/cm*^2^; *p* = 3; *q* = 1; *m*_∞_ = 1/(1 + *exp*(− (*V* + 38)/8.5)); *τ*_m_ = 0.132/(cosh ((*V* + 27)/7.5) + 0.003/(1 + exp (−(*V* + 27)/5))); *h*_∞_ = 1/(1 + *exp*(*V* + 47/6)); *τ*_*h*_ = 10/*cosh*((*V* + 42)/15); *E*_*Na*_ = *RT*/*F* * ln([*Na*^+^]_*o*_/[*Na*^+^]_*i*_); [*Na*^+^]_*o*_ is fixed to be 140 mM; [*Na*^+^]_*i*_ (also written as [Na^+^] in the main text) was dynamically computed by I_Na_ and the Na/K pump; For I_KD_, *g*_*max*_ = 0.216 *S*/*cm*^2^; *p* = 4; *q* = 0; *m*_∞_ = (−0.01 * (*V* + 45.7)/(*exp*(−(*V* + 45.7)/10) – 1))/((−0.01 * (*V* + 45.7)/(*exp*(−(*V* + 45.7)/10) – 1)) + 0.125 * *exp*(−(*V* + 55.7)/80)); *τ*_*m*_ = 1/((−0.01 * (*V* + 45.7)/(*exp*(−(*V* + 45.7)/10) – 1)) + 0.125 * *exp*(−(*V* + 55.7)/80)); *E*_*K*_ = −70 mV; For I_KA_, *g*_*max*_ = 0.02 *S*/*cm*^2^; *p* = 3; *q* = 1; *m*_∞_ = (0.0761 * *exp*((*V* + 94.22)/31.84)/(1 + *exp*((*V* + 1.17)/28.93)))^1/3^; *τ*_*m*_ = 0.3632 + 1.158/(1 + *exp*((*V* + 55.96)/20.12)); *h*_∞_ = 1/(1 + *exp*(*V* + 53.3/14.54)^4^; *τ*_*h*_ = 1.24 + 2.678/(1 + *exp*((*V* + 50)/16.027)); For I_leak_, *g*_*max*_ = 1.25 × 10^−4^ *S*/*cm*^2^; *p* = 0; *q* = 0; *E*_*leak*_ = −65 mV; the Na/K pump model is similar with that used in (17, 40), described by the following kinetic schemes: 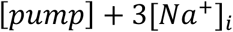 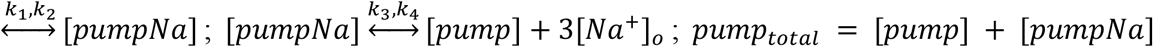; *k*_1_ = 2 *mM*^−3^*msec*^−1^; *k*_2_ = 0.001 *msec*^−1^; *k*_3_ = 0.6 *msec*^−1^; *k*_4_ = 0.437 *mM*^−3^*msec*^−1^; By these parameters, the steady state [Na^+^] is ~ 10 mM when the axon models are silent. For Fig. 4, a scaling factor in the range of 0.2 −1.4 was multiplied with *τ*_*m*_ at all voltages for I_KD_.

For the unmyelinated axon model, all defined currents and the Na/K pump were evenly distributed with C_m_ = 1 μF/cm^2^ and R_a_ = 120 Ωcm. For the myelinated axon model, the ionic currents and the Na/K pump were placed in the nodes of Ranvier (1, 49). For simplicity, we did not model juxtaparanodal regions. At the nodes of Ranvier, C_m_ = 1 μF/cm^2^; On myelinated compartments, there is only a smaller I_leak_ with *g*_*max*_ = 1.25 × 10^−6^ S/cm^2^ and effective C_m_ = 0.01 μF/cm^2^; R_a_ is 120 Ωcm. In both the unmyelinated and the myelinated axon models, the current densities are identical so that we could explore the role of diameter and myelination on Na^+^ accumulation. The *pump*_*total*_ is in the range of 0.5–5 pmol/cm^2^. We did not explicitly simulate the axon initial segment in our model. Instead, we applied strong enough current at the starting end of the unmyelinated model and the first node of Ranvier in the myelinated model to reliably trigger spikes.

## Acknowledgments

The authors would like to thank the constructive suggestions from the reviewers and from the members of Marder lab. This work was supported by the grant R35NS097343.

## Supporting Information

**Figure S1.**
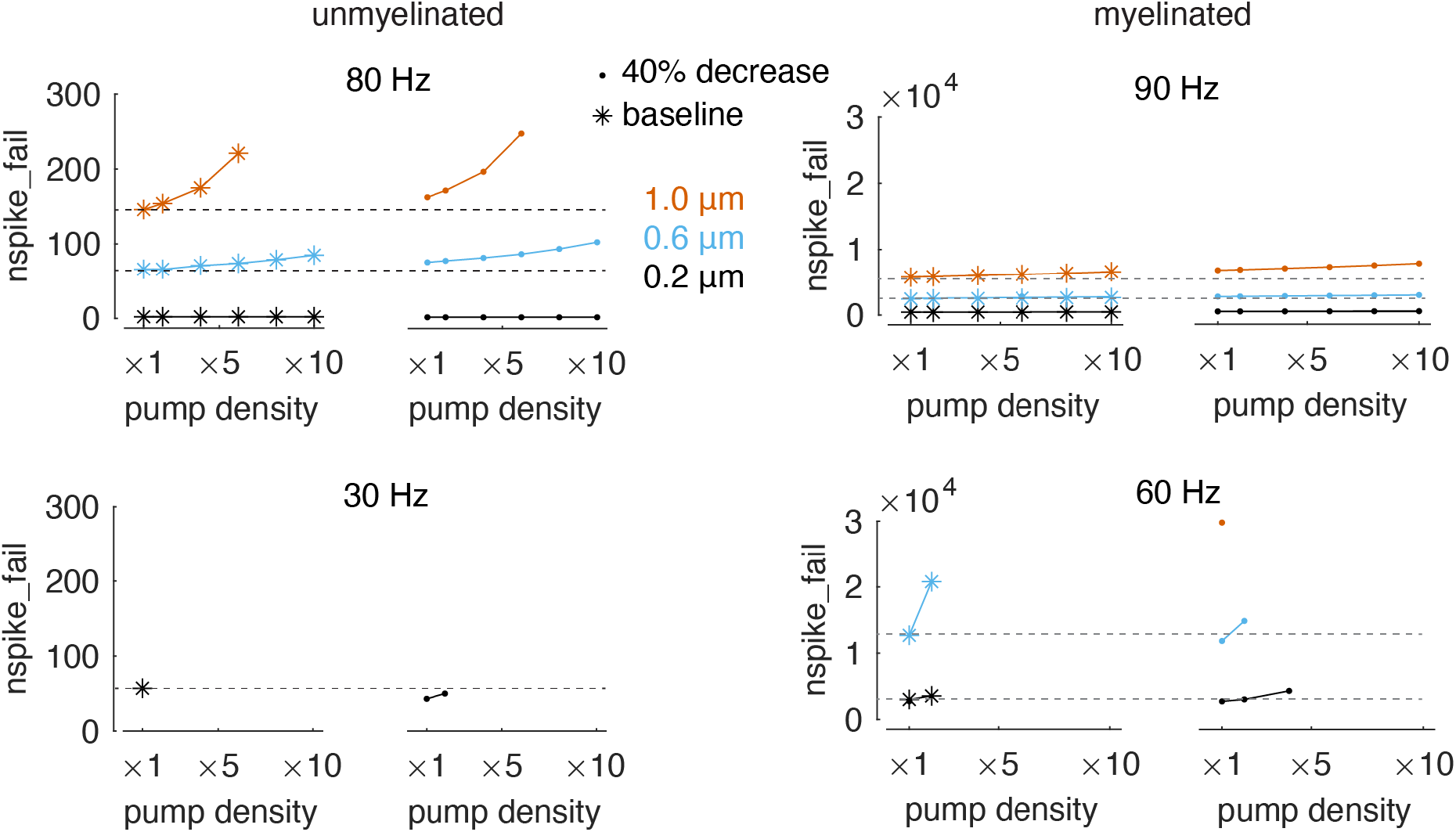
The effects of increasing the rate of Na^+^ current inactivation on propagation resilience. Star represents the baseline model and dot represents 40% decreased inactivation time constant of Na^+^ current. In unmyelinated axon models, at 30 Hz, faster inactivation decreases the resilience in the 0.2-μm-diameter axon model (dots are lower than stars). At 80 Hz, faster inactivation increases the resilience (dots are higher than stars). The same trend is observed in myelinated axon models. The absence of data points means propagation doesn’t fail.

